# An investigation into the critical role of fibre orientation in the ultimate tensile strength and stiffness of human carotid plaque caps

**DOI:** 10.1101/2020.08.25.264457

**Authors:** R.D. Johnston, R.T. Gaul, C. Lally

## Abstract

The development and subsequent rupture of atherosclerotic plaques in human carotid arteries is a major cause of ischemic stroke. Mechanical characterization of atherosclerotic plaques can aid our understanding of this rupture risk. Despite this however, experimental studies on human atherosclerotic carotid plaques, and fibrous plaque caps in particular, are very limited. This study aims to provide further insights into atherosclerotic plaque rupture by mechanically testing human fibrous plaque caps, the region of the atherosclerotic lesion most often attributed the highest risk of rupture. The results obtained highlight the variability in the ultimate tensile stress, strain and stiffness experienced in atherosclerotic plaque caps. By pre-screening all samples using small angle light scattering (SALS) to determine the dominant fibre direction in the tissue, along with supporting histological analysis, this work suggests that the collagen fibre alignment in the circumferential direction plays the most dominant role for determining plaque structural stability. The work presented in this study could provide the basis for new diagnostic approaches to be developed, which non-invasively identify carotid plaques at greatest risk of rupture.

**Graphical Abstract:** 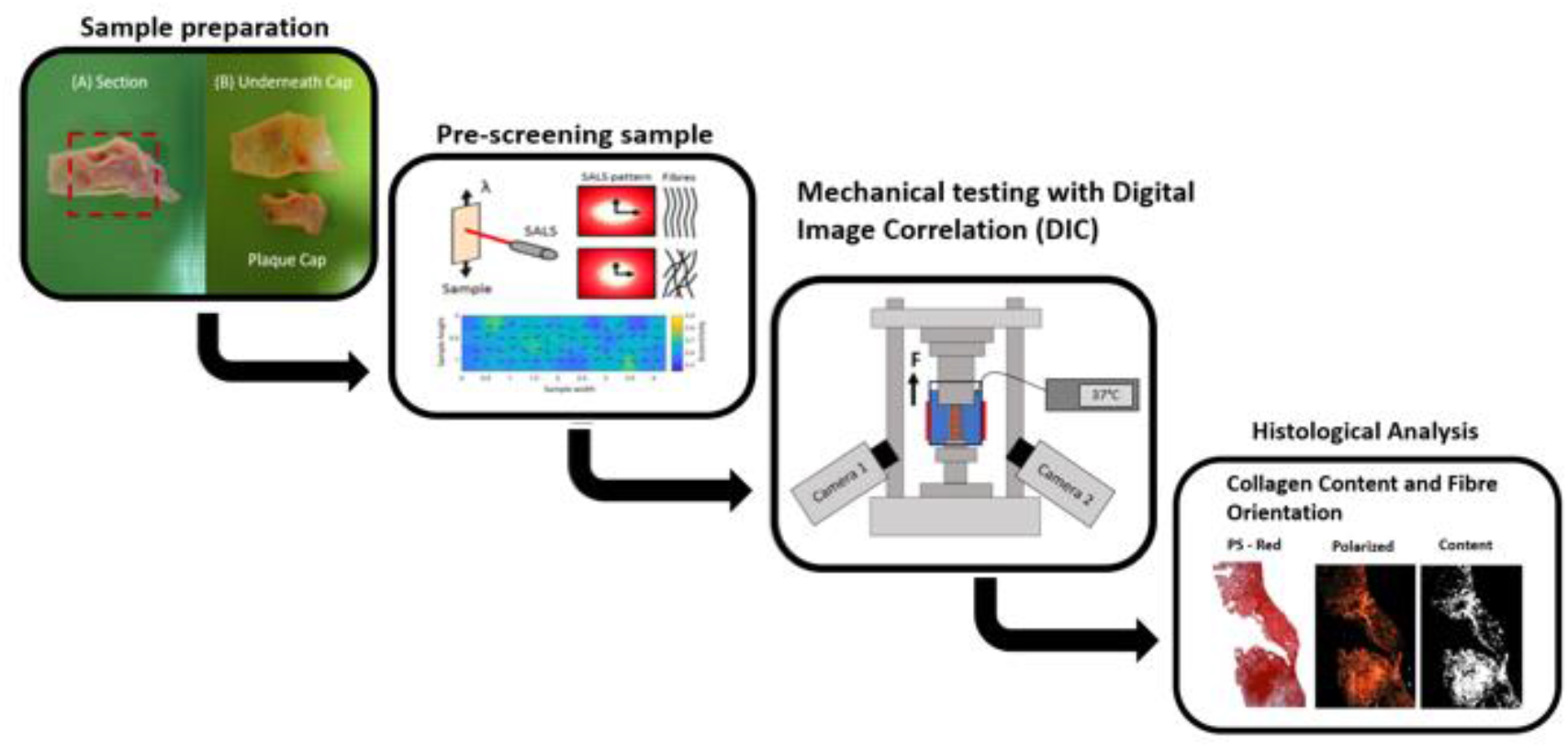

## 1 Introduction

Stroke is known to be one of the leading causes of death and disability in Europe [1,2]. One of the significant contributors to ischemic stroke cases is the development and eventual rupture of atherosclerotic plaques at the carotid bifurcation [3–5]. Atherosclerotic plaques are heterogenous tissues that comprise many different structural components such as lipids, calcium, plaque, haemorrhage and fibrous plaque caps [6], all of which contribute to the overall mechanical response of the tissue. As plaques are subjected to continuous mechanical loading due to blood flow and pressure, failure occurs when the stress exerted on the tissue exceeds its mechanical strength [7]. It is therefore important to understand the underlying structural characteristics of these components and how they dictate the tissue’s overall mechanical response.

It is well known that collagen is the dominant load bearing constituent in arteries [8]. Characterization of the structural integrity of the collagen fibres in atherosclerotic plaques is therefore of utmost importance, since being able to characterize and quantify collagen fibre content and orientation can potentially provide critical insights into the vulnerability of a plaque to rupture [9,10]. There have been some recent advances to quantify the arterial microstructure *in vitro* and *in-vivo* using magnetic resonance imaging [11–15], however, current mainstream *in-vivo* imaging techniques cannot directly image collagen fibres and generally only differentiate between major structural components [16–19]. Therefore, current diagnosis of plaque vulnerability is generally based on geometrical measures such as the degree of vessel stenosis [20–22]. Luminal narrowing is an inadequate metric for atherosclerotic plaque rupture risk, however, as vulnerable plaque rupture is commonly seen below the established clinical stenosis threshold (> 69%) [23]. Consequently, costly and invasive open surgeries, such as carotid endarterectomies, are regularly performed to remove plaques that may be structurally stable, or in cases where better treatment options could be available [24]. There is a need, therefore, for a mechanically sensitive parameter, along with geometric or morphological parameters, which can better identify vulnerable plaques at risk of rupture to better inform treatment planning [25].

Whilst ultrasound studies have suggested that characterizing strains across a plaque can differentiate between stable and vulnerable cases [26,27], these imaging studies do not provide any insights into the underlying characteristics of the structure which determines the tissue’s mechanical behaviour. In a recent *ex-vivo* investigation on porcine carotid arteries, Gaul et al, [2020] showed that collagen degradation in arterial tissue is a strain dependent process, with increased intraluminal pressure leading to higher circumferential strain and higher collagen degradation [28]. That study also showed premature failure of arteries following accelerated collagen degradation due to the significant load bearing role of collagen in the dominant loading direction. Numerical biomechanical models have also identified areas of high stress within the fibrous plaque caps which may be fracture sites in carotid plaque lesions [29– 31]. However, these numerical models do not generally include patient specific plaque properties, and/or collagen content or orientation within the plaques [32], and so the absolute value of these stresses cannot be relied on as a clinical predictor of rupture.

Experimental methods for quantifying patient specific plaque mechanical behaviour have been critical in determining the ultimate tensile strength of these tissues and consequently the risk of plaque rupture [33–35]. Review studies by Walsh et al, 2014 [36] and Akyildiz et al, 2014 [37] summarise the experimental studies that have been performed using uniaxial tests on atherosclerotic plaques and their components, and demonstrate very high mechanical variability across these studies [34,36,38–40]. Davis et al. [2016] focused specifically on the plaque cap, which is often attributed the highest risk of rupture [45]. They found that collagen content was not significant in determining the structural stability of plaque caps, but they also highlighted the need for the fibrous cap collagen architecture to be established in future work [45]. They also noted the critical role that collagenases, such as MMPs, could have in degrading collagen fibres in atherosclerotic plaques, which could lead to a reduction in the tensile strength of the fibrous cap and increased plaque vulnerability [see Davis et al, 2016 cited in 45]. Whilst imaging collagen fibre patterns *in vivo* is still beyond the state-of-the-art in cardiovascular imaging, it is possible to determine such fibre arrangements *ex-vivo,* albeit destructively in the majority of cases such as histology or microscopy [14,47]. In contrast, small angle light scattering (SALS) analysis is a non-destructive technique that can be used to pre-screen fibre structures in thin biological tissue samples *ex-vivo* [48,49] whereby the incident laser light angle is scattered orthogonally to the central axis of the sample’s constituent fibres, providing detail on the sample structure.

Since the role of collagen orientation in the mechanical stability of fibrous plaque caps has not yet been explicitly studied, this study aims to investigate the contribution of collagen fibre orientation and total collagen content to the ultimate tensile (UT) stress, UT strain and stiffness of human carotid fibrous plaque caps. To achieve this goal, the fibre patterns in carotid plaque cap tissues were measured using the non-destructive laser imaging technique, SALS, prior to mechanically testing the samples in uniaxial tension. In addition, robust histological analyses on all test samples were subsequently performed post-testing to further assess collagen fibre orientation and quantify collagen content. To the authors’ knowledge, this is the first empirical assessment of collagen fibre orientation in atherosclerotic plaque caps and its contribution to failure strength and stiffness of this vascular tissue.

## 2 Material and Methods

### 2.1 Sample Preparation

Carotid plaque specimens were obtained from 20 carotid endarterectomy (CEA) patients at St James Hospital Dublin. These symptomatic carotid plaque specimens were firstly washed in phosphate-buffered saline (PBS) to remove residual blood and then stored in tissue-freezing medium (RPMI-60 Media, 1.8 M DMSO; and 0.1 M sucrose), placed into a Mr. Frosty cryosystem containing 2-proponol and cryopreserved at −80°C until the day of testing. On the day of testing, samples were defrosted in PBS at ambient temperature and then sectioned as shown in figure 1. By delineating the cap from the fibrotic media of the plaque, plaque cap samples were dissected to yield circumferential strip samples with a 4:1 (length: width) ratio, as recommended in [36]. Furthermore, dimensions of each plaque cap were recorded (width and thickness) using a light microscope. For accurate analysis of UT stress, UT strain and stiffness, only samples that failed in the centre were included. To prevent sample slippage, velcro was used on the grips.

**Figure 1:**
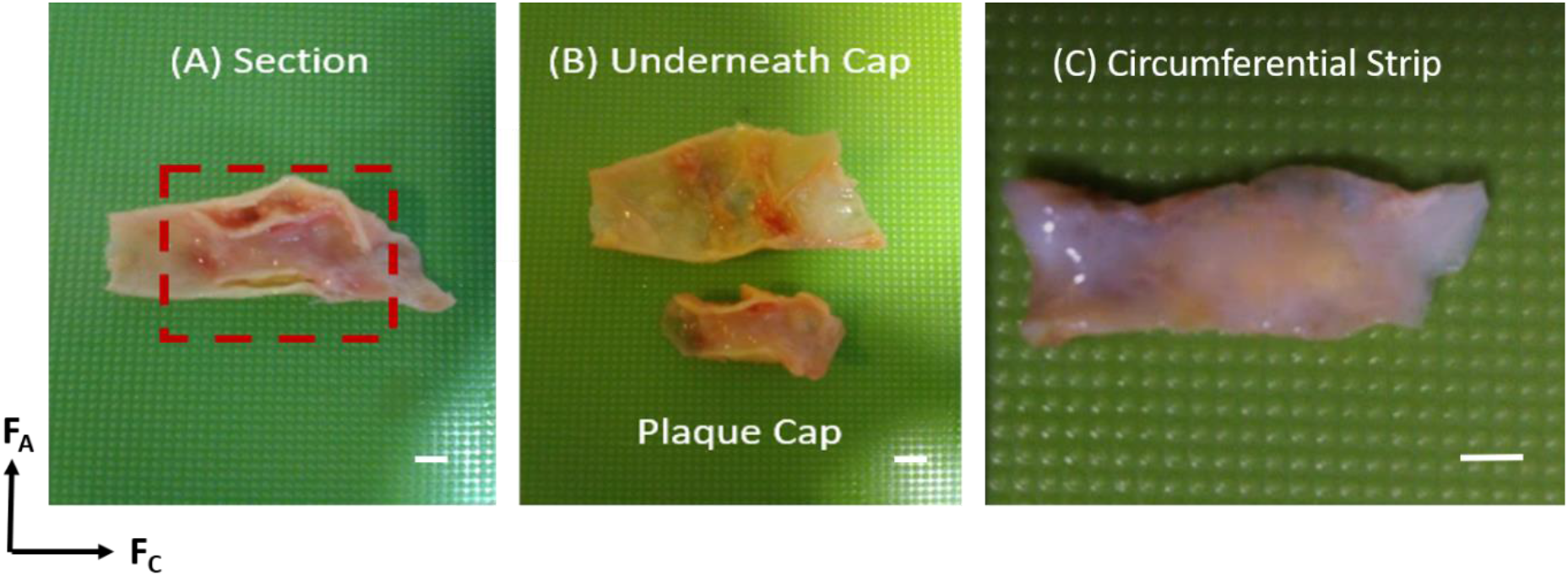
(A) Section of atherosclerotic plaque - Red box indicates the delineation of the fibrous plaque cap from the underlying tissue. (B) Separated plaque cap and underlying tissue (C) Circumferential strip sample of the atherosclerotic plaque cap; scalebar = 1mm.

### 2.2 Small angle light scattering (SALS)

Using an in-house SALS system [49], the dominant fibre orientation of these excised atherosclerotic plaque cap test specimens could be determined before testing. Using a purpose-built MATLAB (MathWorks, Cambridge, UK) code allowing for the pre-dominant direction to be determined, the centroid of the scattered light pattern was established before cycling through all angles from 1 to 360, computing the light intensity at each angle. In SALS, the light is scattered perpendicular to the dominant fibre direction. Raster scanning was performed over a set region of interest and the dominant direction in each region was established. The scattered light profile can either be an elliptical shape if a dominant direction is observed or a circular shape if no dominant direction is observed. This is described by the eccentricity (E), which provides the light distribution at a given interrogation point as a ratio based on the major and minor axes, i.e.

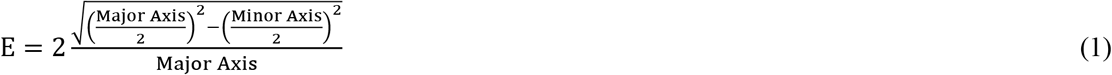

An eccentricity of 1 corresponds to perfect fibre alignment in one dominant direction, while an eccentricity of 0 corresponds to an isotropic distribution of fibres [49], see Figure 2.

**Figure 2:**
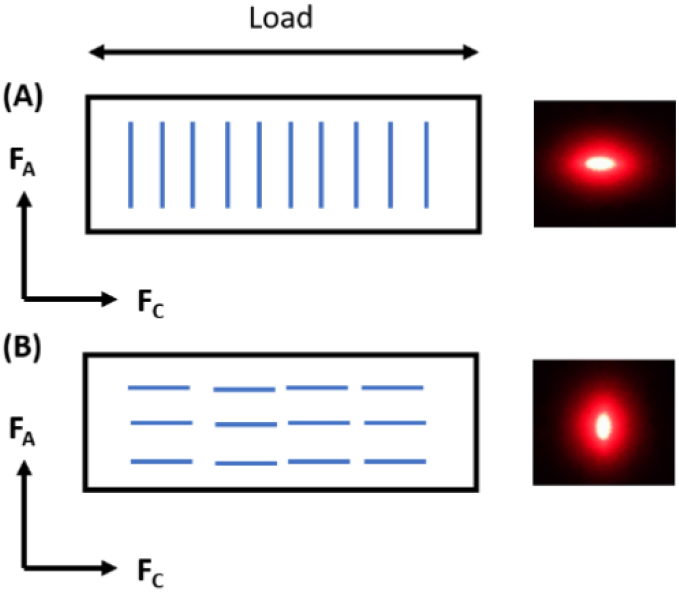
Schematic illustration of the specimen fibre orientation and the resulting scattered ellipse. (A) Fibres in predominantly axial direction (F_A_). (B) Fibres in predominantly circumferential direction (F_C_).

During the pre-screening procedure and due to the variable sample sizes, a 3 mm × 1 mm region of interest was selected in the centre of each sample, thereby allowing a more accurate determination of the dominant fibre direction in the sample in the location of expected failure. After analysis, each sample was then characterized by its dominant fibre orientation, whereby a predominantly axial fibre orientation (F_A_) had fibres perpendicular to the direction of loading (θ from 45° to 90° and −45° to −90°) and a predominately circumferential fibre orientation (F_C_) had fibres parallel to the direction of loading (θ from 0° to ±45°).

### 2.3 DIC and Mechanical Testing

Uniaxial tensile tests to complete failure were performed using a uniaxial test machine (Zwick Z005, Zwick GmbH & Co. Ulm, Germany). All tests were performed in a water bath of PBS solution at 37°C, to simulate the physiological environment. The testing procedure was similar to that outlined in Ghasemi et al, 2018 [50], whereby, a sequence of cyclic loading cycles was imposed on the tissue at a constant displacement rate with strain limits set for each loading step (10%, 20% and 30%). Five preconditioning cycles were set at the beginning of the test to remove viscoelastic effects in the tissue. Digital Image Correlation (DIC) was used to track local strain deformations in the tissue during testing and was set up to record the entire experiment at a rate of two frames per second. A stochastic pattern was applied using spray paint which allowed displacement of the tissue to be tracked, see figure 3. Loading continued in each sample until failure of the tissue occurred and the final force-deformation cycle prior to failure was evaluated to establish the stress-strain behaviour of the samples. Only samples that failed approximately in the centre of the sample were considered for analysis to ensure only robust ultimate tensile stress, strain and stiffness measures were obtained. Ultimate tensile stress and ultimate tensile strain values were then extracted at the point of failure of the sample. Stiffness (E) was also calculated for each sample by taking 10 data points and calculating the slope of the final linear region observed in the stress-strain curves but ending before the final 20% of the curve to ensure consistency across samples [48].

**Figure 3:**
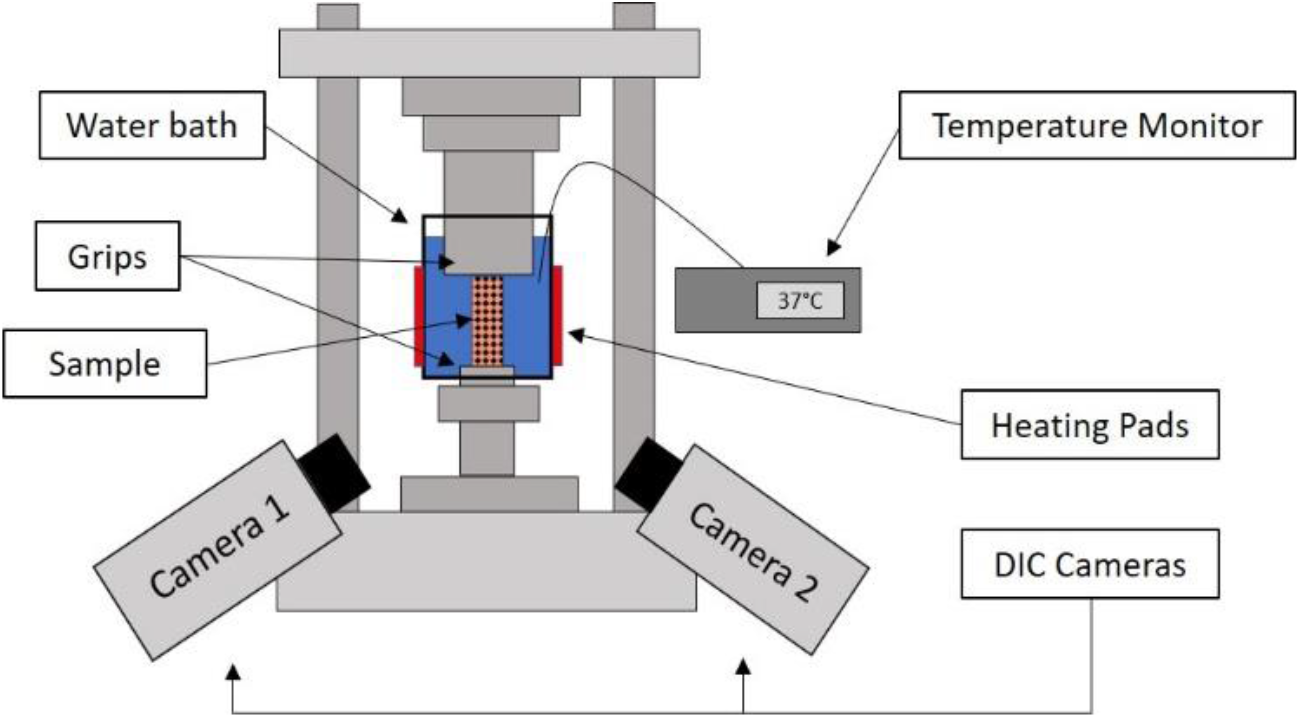
Schematic showing the experimental setup used for the uniaxial tensile tests.

### 2.4 Histology

The use of histology with certain tissue stains such as Haematoxylin and Eosin (H&E) and Picrosirius (PS) red enables quantitative analysis of the tissue microstructure along with microscopy techniques such as polarized light microscopy (PLM) which can extract information on the collagen content and collagen fibre orientation [44–46].

After testing, carotid plaque cap samples were fixed in 10% formalin at 4°C for 24 hours. The samples were then dehydrated (Leica TP1020, Semi-enclosed benchtop tissue processor, Germany) and embedded in paraffin wax blocks. Following this, 8 μm cross sections were cut from the paraffin blocks using C35 microtome blades and floated on distilled water at 37°C before being mounted on glass slides. The slides were left to dry overnight. Staining of the samples was done by using a Leica Autostainer (Leica ST5010, Autostainer XL, Germany), that incorporated the deparaffinization and rehydration of the samples.

Slides were stained with Picro-sirius (PS) red to visualize and quantify the collagen fibre alignment and collagen content [51]. Once stained, the slides were imaged using a brightfield and polarized light microscope (Leica, Wetzlar, Germany) at a range of magnifications (2x and 4x) to allow for detailed analysis. The results for both content and the orientation analysis were determined from an average across two sections taken through the thickness of each sample.

#### 2.4.1 Content Analysis

To determine the content of collagen within tissue samples, post-processing of the PS-red stained tissue sections was performed. Content was determined based on the fractional area of birefringence observed. Firstly, an image of the tissue sample was collected in brightfield and the total tissue area was measured. The same slide was then examined with polarized light and an image was collected and binarized, see Figure 4. This binarization was kept at a constant value (0.2) for all images. The collagen content of the sample was then calculated from comparing the birefringent area to the total tissue area [45].

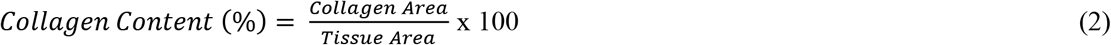

**Figure 4:**
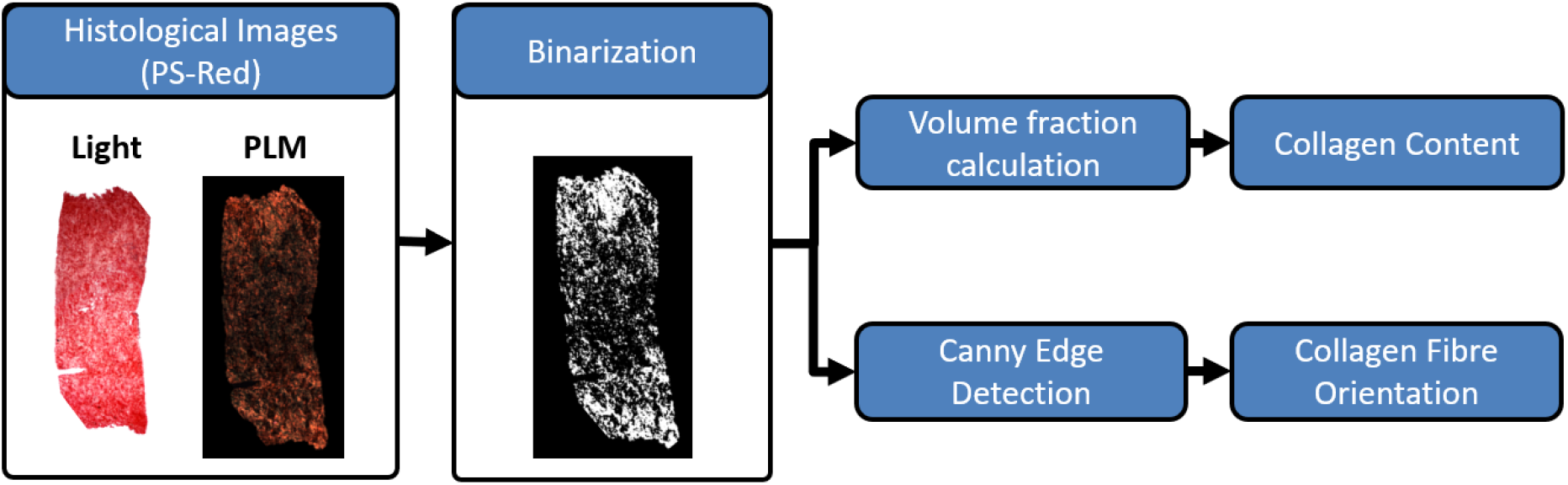
Workflow of processing from histological images to obtain both the collagen content and fibre orientation of atherosclerotic plaque caps.

#### 2.4.2 Orientation Analysis

For collagen fibre orientation assessment via histology, the method used was similar to that reported in Douglas et al, 2016 [44]. A canny edge detection filter was used to isolate the collagen fibres in the image. The filter was applied both horizontally and vertically to remove any noise or speckles in the image and thresholds were set so any small components were removed. Connected adjacent pixels were then found using the *regionprops* function in MATLAB. Delineated fibres containing less than eight or fewer connected pixels were removed, as they were deemed too small to give robust fibre properties [44]. Following this, the code MatFiber [46] was used to determine the mean fibre angle across the sample. The process for content and orientation analysis is summarized in the workflow illustrated in Figure 4.

#### 2.4.3 Statistical Analysis

Statistical analysis was performed with Prism 8 statistical software (GraphPad Software Inc., San Diego, California) and the data for the ultimate tensile (UT) stress, UT strain, stiffness and collagen content were grouped according to the SALS data. Two-tailed unpaired t-tests using Welch’s correction were performed on the UT stress, UT strain, stiffness and collagen content values to investigate statistical significance (i.e. p < 0.05) between the predominately axial fibre datasets and predominately circumferential fibre datasets due to the uneven sample sizes in each group.

## 3 Results

### 3.1 Thickness measurements across atherosclerotic plaques

Three thickness measurements were recorded for all atherosclerotic plaque cap samples that underwent testing and are recorded in Figure 5. Plotted as mean and standard deviation for each, every individual point corresponds to one measurement. Overall, the thickness of our samples is in the range 0.5398 ± 0.114599, showing the thickness of the plaque cap varies within and between samples, and that this can be attributed to the heterogeneity of the tissue.

**Figure 5:**
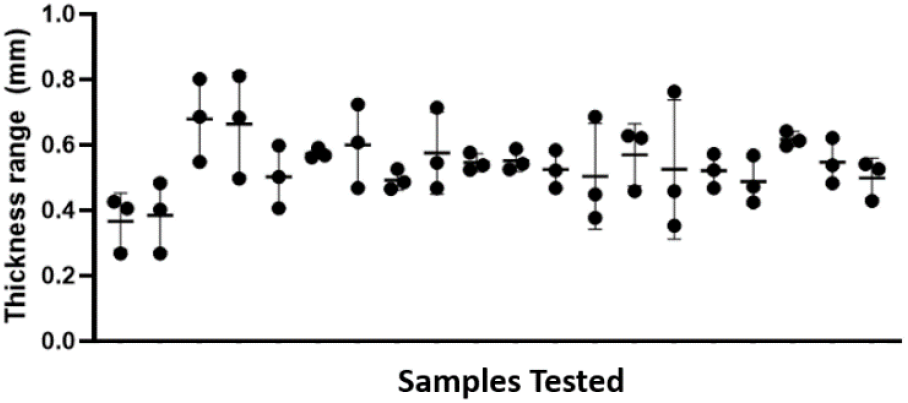
Thickness measurements taken across the atherosclerotic plaque cap samples tested. Plot details the mean and standard deviation of the measurements taken.

### 3.2 Dominant Collagen Fibre Orientation in Carotid Plaque Caps using SALS

The mean fibre angle (θ) and standard deviation within samples was determined using SALS and is shown in the order in which samples were tested in Figure 6, where considerable variation in the dominant fibre angle is evident across these plaques harvested from different patients. The samples were therefore grouped in terms of their dominant fibre direction, whereby, predominantly axial fibre orientation (F_A_) samples are those where fibre angles, θ from ±45° to ±90°, and predominantly circumferential fibre orientation (F_C_) samples have fibre angles, θ from 0° to ±45°. After determining the respective groupings, Figure 7A, 7B show both the SALS eccentricity plots and fibre (azimuth) angle distributions for all samples, detailing the sample to sample variation in eccentricity values and fibre distributions. The azimuth angle is an angular measurement in a spherical coordinate system whereby, for our analysis, an azimuth angle at 0° corresponds to the circumferential direction and ±90° corresponds to the axial direction.

**Figure 6:**
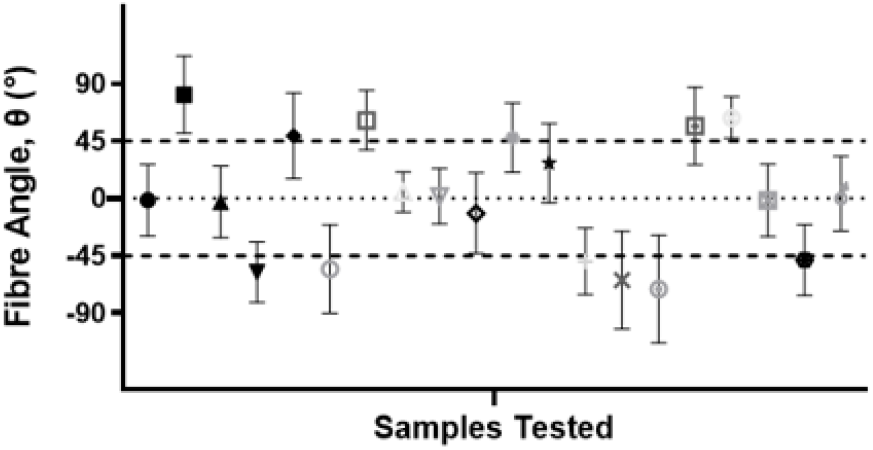
Mean Fibre Angle, θ, and standard deviation obtained from samples in the order of testing. Samples where θ is from ±45° to ±90° are denoted as samples with predominantly axial fibres (F_A_) and those with, θ from 0° to ±45° are denoted as having predominantly circumferential fibres (F_C_). Each datapoint is one sample.

**Figure 7:**
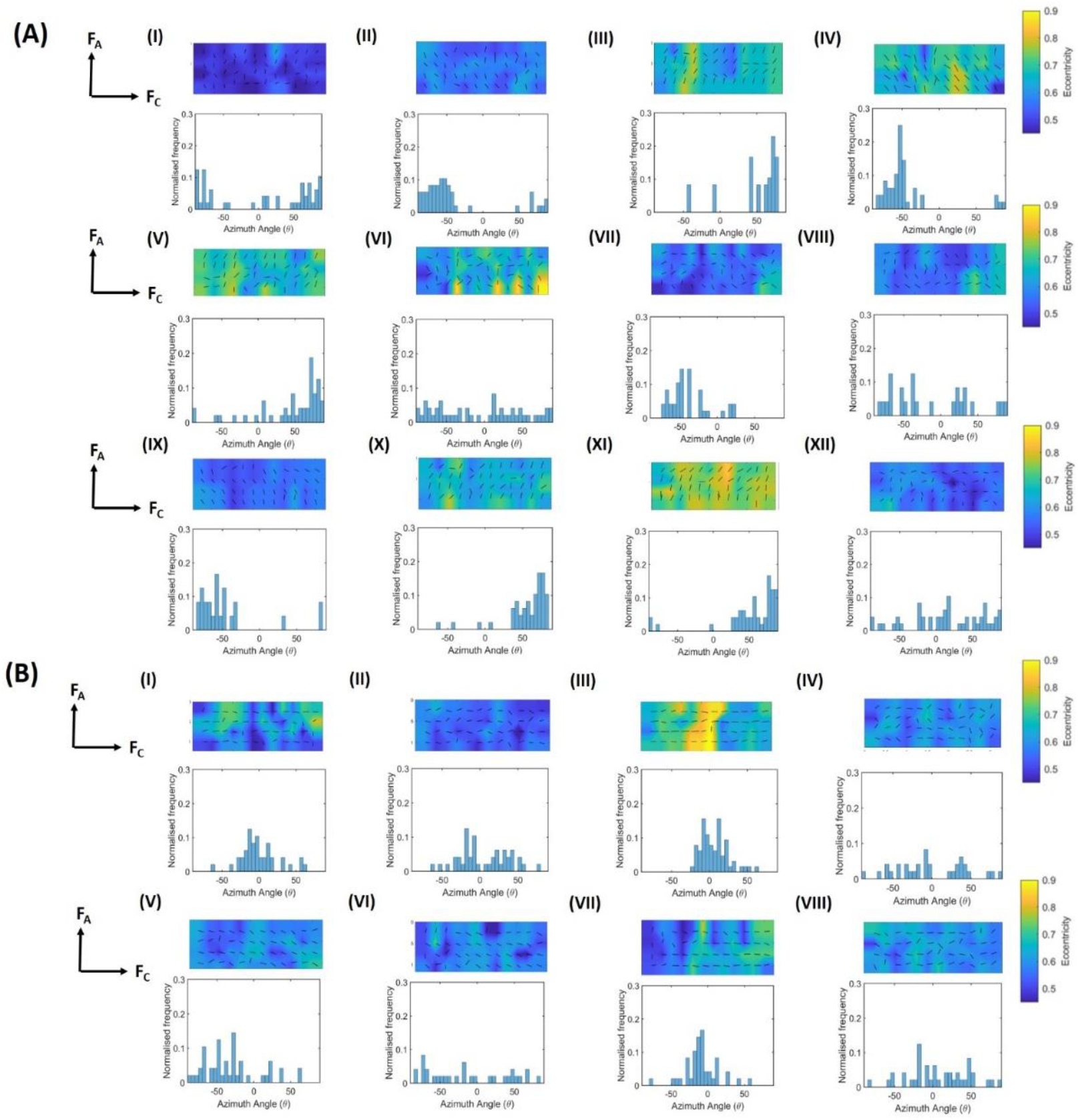
SALS eccentricity plots and histograms displaying fibre angle distributions after grouping into (A) predominantly axial fibre dataset (F_A_) and (B) predominantly circumferential fibre dataset (F_C_).

### 3.2 Ultimate Tensile Strength of Carotid Plaque Caps

Data presented here are from samples that failed in the centre of the tissue sample, with a representative sample shown in Figure 8.

**Figure 8:**
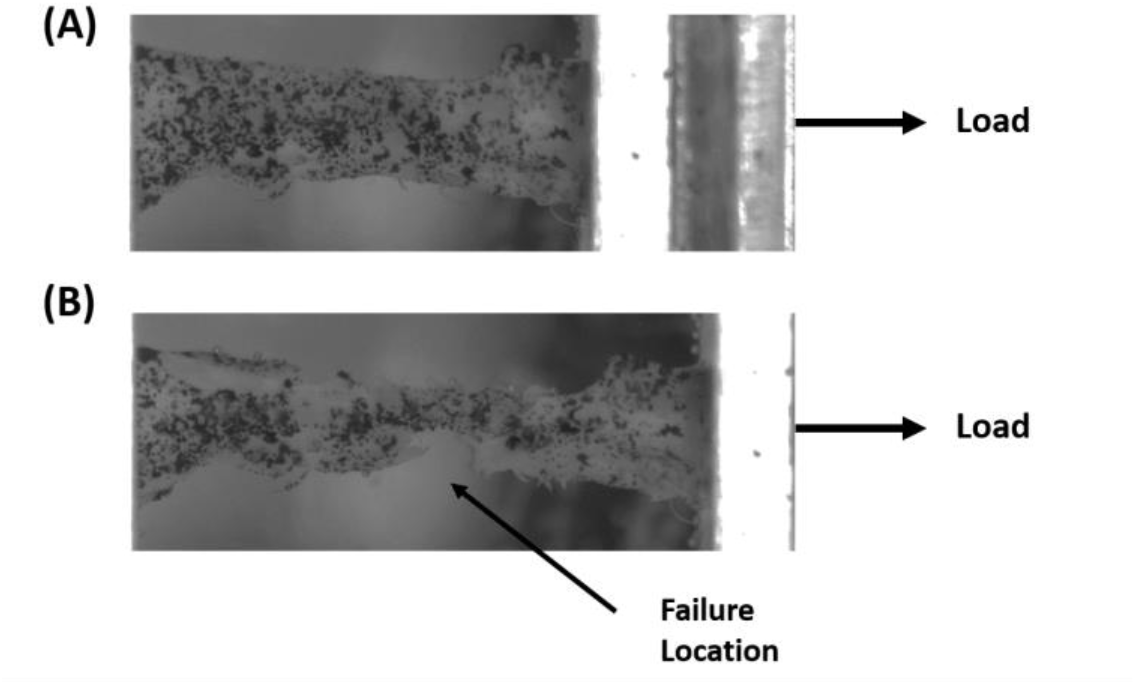
DIC view of plaque cap with speckle pattern undergoing uniaxial test showing (A) initial sample setup and (B) failure at centre of sample.

Stress-strain curves for all samples demonstrated the characteristic non-linear J-shaped curves typical of vascular tissue (Figure 9A-B). To observe the effect of fibre orientation, the uniaxial test results were again grouped according to the predominant fibre direction, namely axial fibres and circumferential fibres based on the SALS results, see Figure 9A and 9B, respectively.

**Figure 9:**
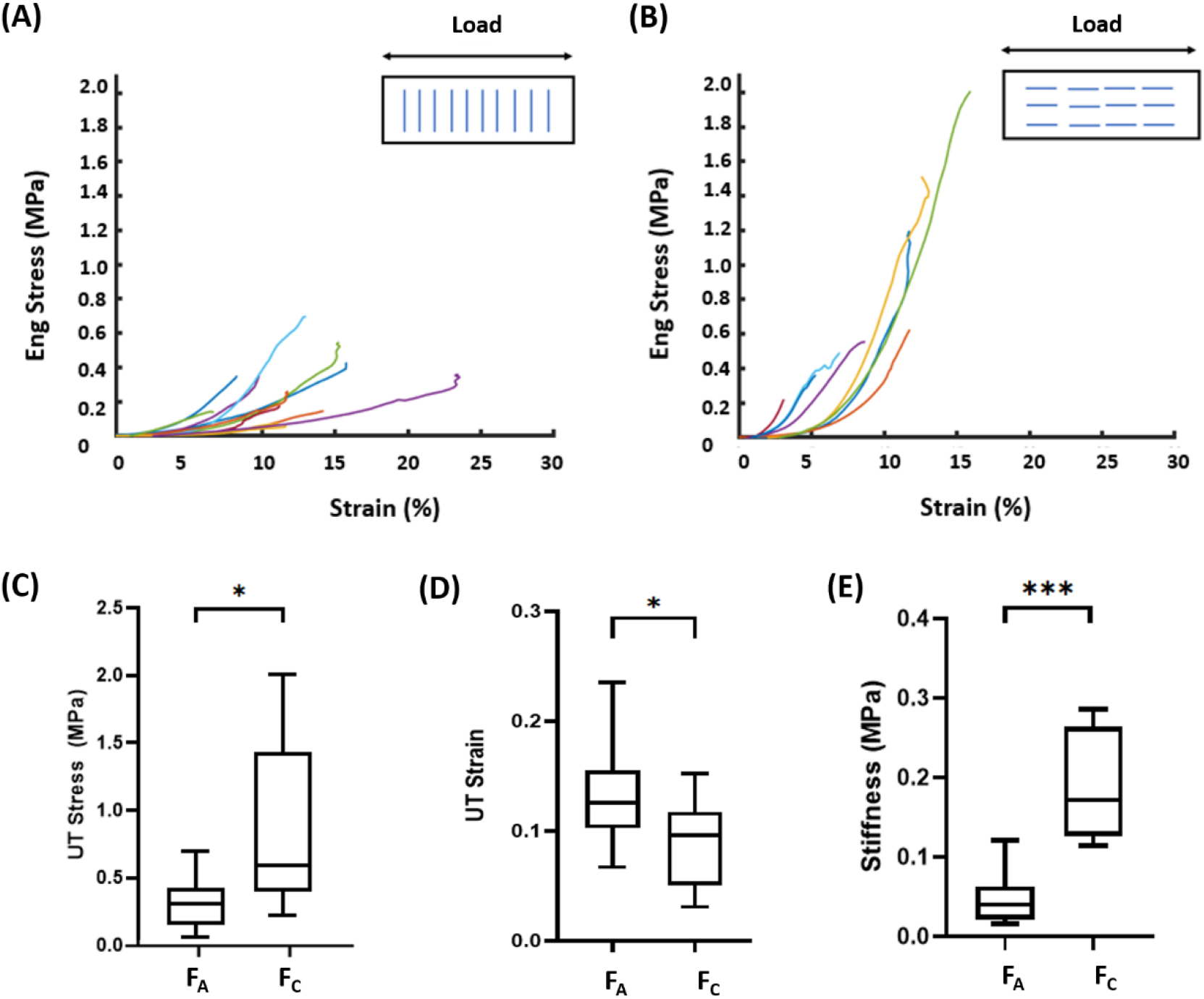
Engineering stress-strain curves for A) predominantly axial fibre datasets (F_A_) and (B) predominantly circumferential fibre datasets (F_C_). Statistical analysis detailing the significant difference between (C) Ultimate tensile stress, (D) Ultimate tensile strain and (E) Stiffness between the groupings, * p < 0.05, *** p < 0.0005.

Figure 9C, 9D and 9E demonstrate the significantly different results between the samples with predominantly axial and circumferential fibres for UT stress, strain and stiffness. The mean UT strain is significantly higher (0.1315 ± 0.0434 versus 0.0961 ± 0.04085), and mean UT stress significantly lower (0.3068 ± 0.1827 versus 0.8718 ± 0.6322) in the predominantly axial fibre datasets, whilst the mean stiffness of samples with predominantly axial fibres is also considerably lower than the samples with predominantly circumferential fibres (0.04547 ± 0.03092 versus 0.1869 ± 0.06785), see figure 9E.

### 3.3 Histological Analysis of Carotid Plaque Caps

#### 3.3.1 Histological Evaluation of Collagen Fibre Orientation and Content in Atherosclerotic Plaque Cap Sample

Collagen content for samples with fibres predominantly in the axial direction was found to have a wider range of content values when compared to samples with predominantly circumferential fibres, see figure 10B. No significant difference in collagen content was identified between the datasets, however, suggesting that collagen content does not play a significant role in determining the UT stress, UT strain and stiffness of the tissue.

**Figure 10:**
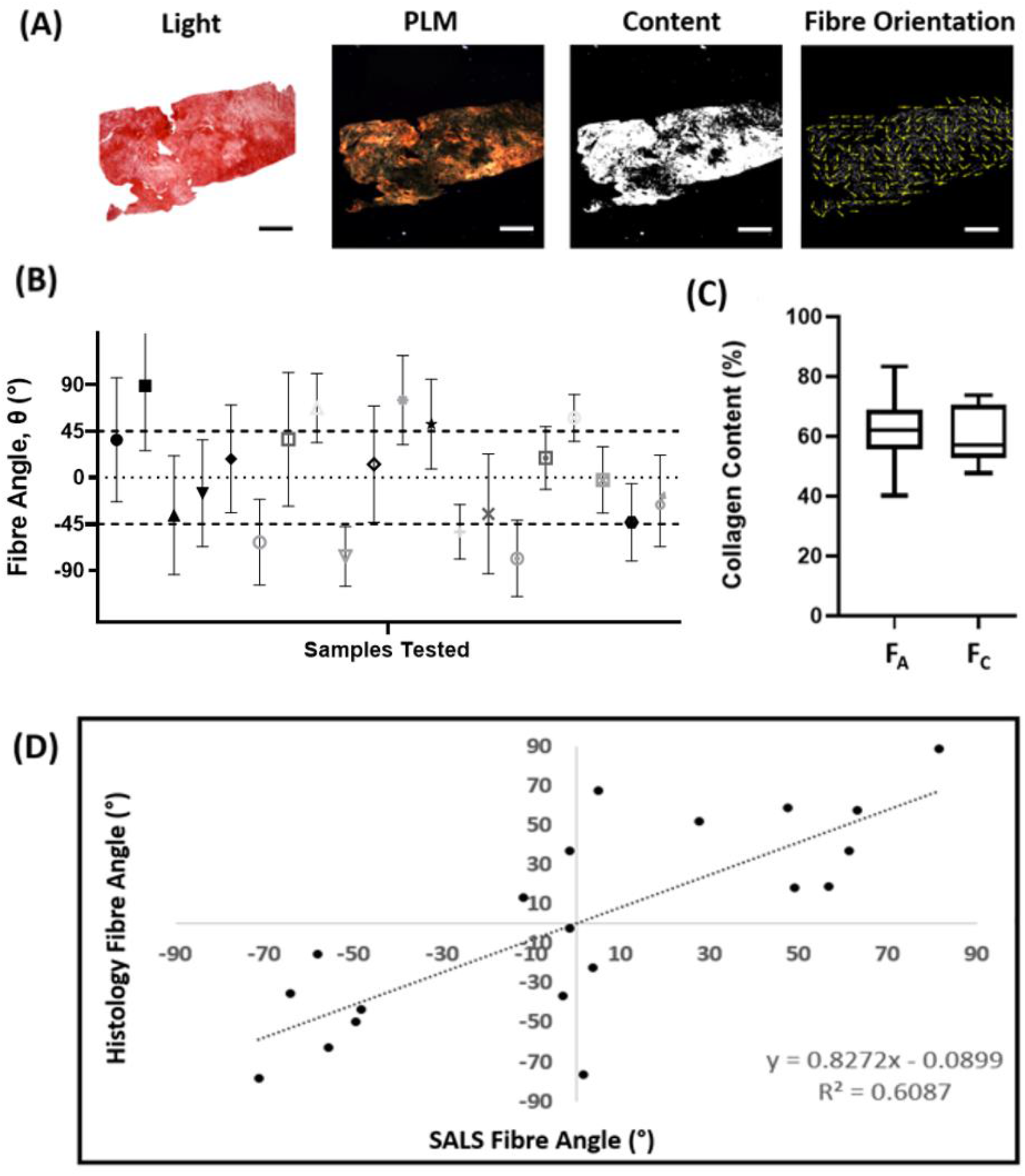
(A) Representative images detailing the post-processing of tested plaque cap tissue after histological staining with Picrosirius red; scalebar = 1mm. (B) Mean fibre angle and standard deviation extracted from samples in the order of testing. (C) Collagen content comparing the predominately axial fibre group and predominately circumferential group. (D) Pearson correlation test between SALS fibre angle and calculated fibre angle from histology.

From our results, it is shown that the histological fibre angle obtained is quite different to the SALS fibre angles shown in figure 6. A Pearson correlation analysis performed between the two datasets showed a relatively weak correlation between the SALS fibre angle measurements and the histological fibre angle calculated, with an R^2^ = 0.6087, see figure 10D. Whilst SALS obtains an average fibre angle from the laser light passing through the tissue prior to testing, histology is performed on only a limited number of thin sections of the tissue and it is also performed after mechanical testing. In addition, histology requires a number of processing steps before final analysis which can induce sample shrinkage. The process also relies on obtaining a flat cross section of tissue during sectioning to ensure robust fibre angle calculations.

Figure 11A and 11B show a poor correlation between collagen content and UT stress (R^2^ = −0.0192) and UT strain (R^2^ = 0.0227), respectively. These findings are independent of fibre orientation.

**Figure 11:**
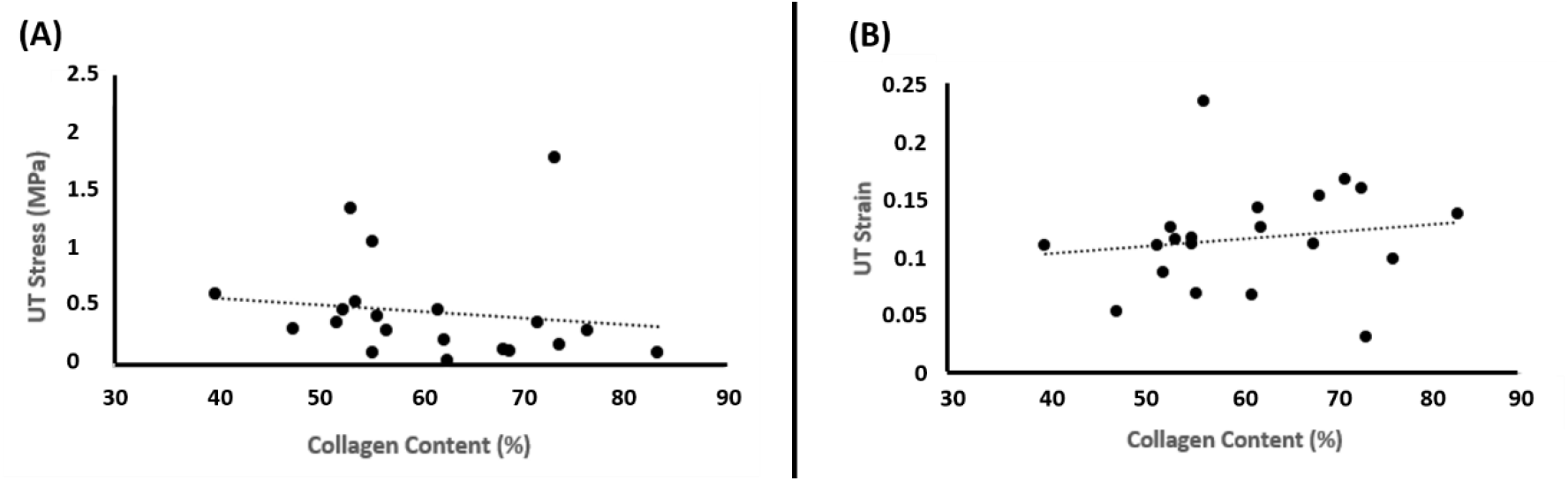
Correlation plots for (A) collagen content and ultimate tensile stress and (B) collagen content and ultimate tensile strain, when not considering fibre angle groupings.

## 4 Discussion

The highly variable nature of the mechanical properties of atherosclerotic plaques has been observed in our samples, validating the observations seen in previous studies [34,36,38–40,45]. Throughout the study, SALS was used as a pre-screening tool to determine the dominant fibre orientation of our samples, thereby determining the anisotropic nature of the atherosclerotic plaque cap. The eccentricity in our samples was low at approximately 0.6 (Figure 7) which would suggest that whilst the cap has an anisotropic collagen structure, it may not be coherently aligned in a particular direction. Thickness variability in the samples can also contribute to this low SALS eccentricity measure. Despite the eccentricity values, once samples were grouped based on SALS fibre angle measurements into those with predominantly axial fibres and predominantly circumferential fibres, our results demonstrate that the orientation of the collagen fibres in the plaque caps plays a significant role in determining the ultimate tensile strength and strain in the plaque caps. The plaque caps with predominantly circumferential fibres demonstrate superior load bearing capacity with higher stresses borne before failure when compared to the predominantly axially aligned fibre samples. Interestingly, our histological results suggest that collagen content alone does not play a dominant role in the strength and stability of atherosclerotic plaque cap tissue given that no significant difference was observed in collagen content between the two plaque cap groupings. Furthermore, when we look at the correlation of collagen content to the ultimate tensile stress and strain, independent of groupings, we do not see a correlation between the collagen content and ultimate tensile stress and strain, suggesting that collagen content alone does not play a significant a role in the load bearing capacity of the tissue.

A particularly interesting finding from this work is the failure of samples with fibres in predominantly the axial direction at lower UT stress and higher strains than the samples with a predominantly circumferential fibre arrangement. This key finding offers potential mechanistic insight into the results demonstrated in previous *in vivo* imaging studies such as Huang et al, 2016 [27], where vulnerable plaques were found to have a higher maximum value of absolute strain rate from diastole to systole. Without knowing the mechanical structure of the plaque and vessels in the Huang et al, 2016 [27] study, our results would suggest that these vulnerable plaques could possibly have fibres more axially aligned which would consequently fail at lower UTS and thereby make them more vulnerable to rupture. This hypothesis clearly warrants further investigation in *in vivo* imaging studies on potentially vulnerable plaques, but it could possibly open up new diagnostic techniques with a clear mechanical origin.

Finally, as outlined in Davis et al, 2016 [45], the use of stress as a rupture metric, as used in many computational studies [29,30], may not be the most appropriate mechanical measure to use when trying to determine plaque rupture vulnerability. This is since plaque caps with collagen oriented in the dominant load bearing direction can withstand higher stresses, however, without this microstructural insight, the stress alone does not offer a robust rupture index, especially as our study highlights that collagen content alone does not correlate with the UT stress and UT strain. Stress can only act as a true rupture measure when it can be determined accurately and compared to a known tissue rupture strength. In contrast, tissue strain measures may offer insights into the underlying microstructure, particularly when considered in the context of the data presented here where higher strains are consistently associated with tissue of lower fracture strength. Characterizing locations across a plaque cap where the deformation of the tissue is increased compared to normal arterial deformation values would suggest microstructural changes to the tissue and overall structural weakness. Furthermore, the current study addresses the limitations stated in Davis et al, 2016 [45] and includes the fracture behaviour of fibrous plaques with respect to their underlying fibre orientation, finding that plaque caps with axially aligned fibres would strain more and fail at lower UT stress.

Whilst this study provides critical new insights into the mechanical behaviour of atherosclerotic plaque caps, there are some limitations in the study; firstly, only fibrous plaque caps were tested and other components that could contribute to the overall mechanical response of the plaque were not considered here. Calcifications can lead to localized stress concentrations and can even potentially alter the fibre orientation within the plaque cap [52] and would therefore be of interest in future investigations. Furthermore, the influence of macrophages was not investigated in this study. Macrophages are known to infiltrate the plaque cap during inflammation and can potentially degrade the matrix of the tissue, thereby weakening the mechanical integrity of the plaque [53]. Whilst macrophage content was not directly explored, the influence of matrix degradation was investigated by quantifying collagen content within the plaques.

## 5 Conclusion

Carotid plaque cap rupture is a local mechanical event in the vessel wall, whereby the stress exerted on the plaque exceeds its mechanical strength causing it to rupture. Parameters such as the percent stenosis do not provide robust metrics for determining plaque vulnerability especially for lower grades of stenosis as they do not consider the underlying mechanical strength of the plaque tissue. The current study demonstrates that the mechanical integrity of the plaque cap is governed by collagen fibres and that collagen content alone may not be a robust predictor of plaque rupture. The critical role of collagen fibre orientation relative to the dominant loading direction in the vessel, shows that for maximum strength, collagen fibres should be in the load bearing circumferential direction. When collagen fibres are predominantly in the circumferential direction, the tissue exhibits a higher ultimate tensile strength and overall stiffer behaviour with lower strains, as compared to plaque tissue with collagen fibres oriented in a predominantly axial direction. The is important as being able to characterize the strain using *in-vivo* imaging techniques, could potentially aid in the identification of fibre orientations and the overall mechanical strength of the tissue and thereby offer a more mechanistic basis for a clinical indicator of carotid plaque rupture risk.

## Conflicting Interests

There is no conflict of interest to be declared by the authors.

## Funding

Research was supported by the European Research Council (ERC) under the European Union’s Horizon 2020 research innovation programme (Grant Agreement No. 637674).

## Ethical Approval

Ethical approval for obtaining the plaques in this study was obtained from the St James Hospital ethical committee in compliance with the declaration of Helsinki.

## Acknowledgements

The authors would like to acknowledge the vascular department in St James Hospital. Dr Prakash Madhavan, Dr. Zenia Martin, Dr. Adrian O’Callaghan and Dr. Sean O’Neill. A special thanks goes to vascular registrar’s Dr Lucian Iacob, Dr. Mike Bourke and Dr. Aoife Kiernan for consenting patients in this study.

## Notes

### Competing Interest Statement

The authors have declared no competing interest.

